# Designing an intuitive web application for drug discovery scientists

**DOI:** 10.1101/169193

**Authors:** Nikiforos Karamanis, Denise Carvalho-Silva, Jennifer A. Cham, Luca Fumis, Samiul Hasan, David Hulcoop, Gautier Koscielny, Michael Maguire, William Newell, ChuangKee Ong, Eliseo Papa, Andrea Pierleoni, Miguel Pignatelli, Sangya Pundir, Francis Rowland, Jessica Vamathevan, Xavier Watkins, Jeffrey C. Barrett, Ian Dunham

## Abstract

Although a scientific web application that is intuitive can help scientists utilize data more easily and advance their research, there is little guidance on how to design such an application in the academic literature. We discuss how we designed an intuitive application for bench scientists working in drug discovery following an approach that can be applied to the design and development of scientific resources in a broad range of disciplines.

## Motivation

In a recent commentary, Ascoli *et al.*^1^ encouraged resource developers to “design and implement intuitive ergonomics” so that scientists become more keen to share data in public repositories. Previous work in bioinformatics has pointed out the need to improve the usability of resources for non-bioinformaticians^2,3,4,5,6^. More generally, a scientific web application that is user friendly can help scientists utilize data more easily and advance their research.

However, there are very few case studies available in the academic literature to provide guidance on how to develop a user friendly web application for wet lab scientists. To fill this gap, we describe here how we developed the Open Targets Platform (www.targetvalidation.org) specifically focusing on an intuitive design.

The Open Targets Platform supports bench scientists working on drug discovery in academia and the pharmaceutical industry to identify and prioritize drug targets faster and with more confidence. We applied Lean User Experience (UX) design^7^ methods to understand and address the needs of our users. This involves engaging with users from the very earliest stages of design and including them in collaborative design and evaluation activities throughout development.

Despite the specialized audience of our Platform, our approach can be applied to the design and development of other web applications made for scientists irrespective of their discipline. We hope that this paper will inform and inspire other members of the bioinformatics community wishing to design and develop intuitive web resources for lab scientists.

## User experience design

UX design (which is often referred to as User-centered Design, UCD) focuses on understanding the needs of users and ensuring that these needs are met^8^. The developers of an application often believe that users will find the application as easy to use as they do but this assumption turns out to be false most of the time^9^. This difference in perspective is not uncommon in the development of scientific web applications. For example, wet lab scientists often find bioinformatics applications hard to use^2,3,4,6^. One of the main goals of UX design is to help those responsible for a service move away from the “self-as-user” outlook and empathize more with their users in order to develop more user friendly applications.

Lean UX design^7^ stems from recent Lean and Agile development practices with short cycles of iteration. Lean UX design tries to address the main challenges in developing an intuitive application in an Agile context: Like traditional UX design, Lean UX design places the needs of users at the center of the development process. Additionally, it advocates close collaboration between the members of the development team and engaging with users early and often in the design process.

Although Lean UX design has been applied to the production of a wide range of digital services and products, we are not aware of another case study in the academic literature where it has been applied to the design of a bioinformatics web application for bench scientists. de Matos *et al.*^10^ present a case study in applying UCD to enhance a non-iterative approach for developing a bioinformatics application. We extend their work by emphasizing the importance of collaboration within an iterative development process and by discussing how we supplement qualitative user feedback with quantitative metrics, which were also defined collaboratively.

## Fostering collaboration

The Open Targets Platform integrates different types of biological data from several public databases^11^. Our aim was to design the Platform so that drug discovery scientists can use it without requiring in-depth familiarity with the data integration methods or expertise in bioinformatics. To achieve that we had to approach the Platform from the perspective of our users.

In addition to bioinformaticians, our multidisciplinary team consists of software developers, academics and researchers employed by industrial partners. As in many other scientific projects, we needed to work with each other to succeed.

Our team also includes a UX designer who prepares user research, problem definition as well as design and testing activities in close collaboration with the other members of the team and facilitates these activities making sure that they include both developers and users. Weekly meetings ensure the progress of development is in line with the agreed UX design. We also turned our workspace into a design studio with diagrams and designs on the walls to foster a sense of shared ownership and enable discussions focused on the needs of our users.

Supplementary Figure 1 illustrates how a drug discovery scientist can use the Platform to answer her key questions. In the rest of this paper, we discuss in more detail the Lean UX methods that we used to engage with our users and to work with each other in designing the Platform.

## Empathizing with users

We embarked on the design process by interviewing 27 scientists and managers working on drug discovery for various academic and industrial organizations. The interviews were carried out by several team members and we developed an interview guide covering a wide range of issues related to drug target identification to ensure consistency.

Each interviewer captured the main insights of the interview in a systematic way using an empathy map^12^ (see Supplementary Fig. 2). The empathy maps were shared with the rest of the team and used to create diagrammatic representations.

We also observed scientists working in early stages of target identification. This gave us the opportunity to witness the complex activities that were mentioned in the interviews in a real setting. The observed activities were analyzed and visualized as diagrams in interpretation sessions with the rest of the team^13^ (see Supplementary Fig. 3). Researching and analyzing users’ activities in these ways helped us empathize with them.

## Focusing on user needs

User research generated a lot of information, which needed to be synthesized to identify the most important user needs that the Platform should address. To achieve this we tried to answer the following questions based on the information that we collected:

### a) Who are our users?

We created personas^7,9^ to represent the various stakeholders in drug target identification and prioritization, and to focus our efforts on the most important user needs. “Pat” in Figure 1 represents our main user group, wet lab scientists working on drug discovery. The information that Pat needs to answer her research questions is dispersed in different resources and not easily accessible to her. Support by a bioinformatician is limited in supply, even for simple queries related to Pat’s research.

**Figure 1:**
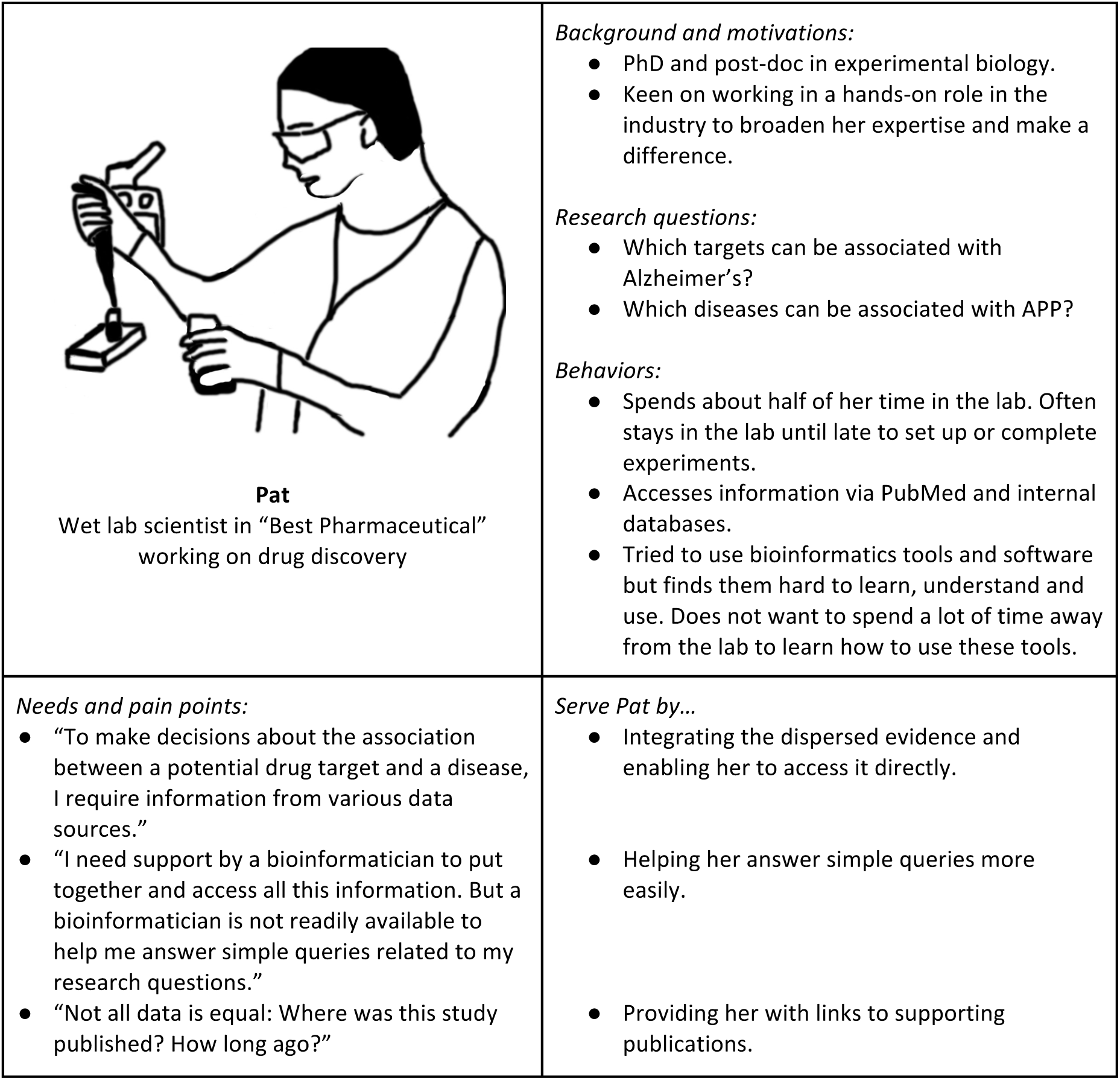
Pat represents our main user group, wet lab scientists working on drug discovery. Creating Pat’s persona enabled us to concentrate our efforts on what matters most for our users.

### b) What problem are we trying to solve?

Identifying Pat’s needs and pain points helped us to define the main problem that we were trying to solve with our Platform. Primarily, our users have difficulty answering the following questions:

- Starting from a particular disease (e.g. Alzheimer’s), which targets can be associated with this disease?
- Starting from a particular target (e.g. APP), which diseases can be associated with this target?

Therefore, we focused the design of the Platform on providing scientists like Pat with more direct access to the evidence that can be used to answer her two main questions. The aim of the Platform is not to answer the questions on behalf of or instead of Pat. Rather, we want to allow Pat find the available evidence so that she can use her expertise in target identification to decide which associations are the most promising to pursue.

### c) How can we improve on current work practice?

Although the two questions in our problem definition look simple, the work practices involved in answering these questions can be quite complex. In order to understand what is typically involved in answering these questions, we created diagrammatic scenarios which synthesized information from the interviews and observations. We characterized the scenarios in terms of the data that people use, the actions that they perform and the inputs and outputs of these actions (see Supplementary Fig. 4).

To capture the wider context in which the scenarios take place, we collaboratively created a diagram of the knowledge that we gained through our user research (see Supplementary Fig. 5). This diagram (which we referred to as the “target identification landscape”) represents the common themes in early target identification and prioritization that we identified. The landscape diagram and the scenarios were placed on the wall of our studio so that we could refer to them throughout the design process.

We concluded the definition stage by specifying the scope of the Platform (see Supplementary Fig. 6). This chart was used to communicate the main proposed feature concepts of the Platform to the Open Targets leadership team. The feature concepts were organized in three tiers, each corresponding to a distinct benefit for Pat and the user group that she represents. In producing this chart, we also started to think about what would constitute success for the Platform (see section about metrics below for more details).

Our approach is lean because we focus on understanding users’ needs before writing extensive documentation or code. However, the various artefacts that we use as communication and collaboration tools (see Fig. 1, Supplementary Fig. 2, Supplementary Fig. 3, Supplementary Fig. 4, Supplementary Fig. 5 and Supplementary Fig. 6) are very rich in information, helping us move away from the ‘self-as-user’ outlook.

## Working iteratively

User research helped us understand whom we want to help, the problems that we want to solve for them, how our users currently go about solving these problems and how we can improve on current work practice with our Platform.

Co-creating this knowledge established common ground for future development work within our team. Involving users and their managers as the knowledge sources generated and maintained their interest in the Platform. Sharing our main insights with the leadership team made them trust us and buy into our vision. It is important to emphasize that all this happened before we started developing the web application.

To develop the Platform we next began working in small iterations, each lasting two to three weeks. Each iteration consisted of a design and prototyping phase which was followed by testing and review activities. We delivered the first Beta version of the Platform (Beta v01) after only 7 iterations from the initial designs on paper (see Supplementary Fig. 7).

## Diversity and alignment through design

We approached the design phase of each iteration as a collaborative task and ran a series of design workshops with our development team. For each workshop we prepared a statement of the problem we were trying to solve and collected samples of data related to this problem.

For example, we ran an internal workshop to sketch a visual overview of the quality of the various types of data that support a target-disease association. We explored a range of solutions (see Supplementary Fig. 8) and shortlisted a few options to show to users for feedback. This included an early version of the evidence visualization (the “flower”) that we use in the Platform (see Supplementary Fig. 1d).

Sketching out ideas on paper is a quick and inexpensive way to explore the design space^14^. Collaborative sketching has the added benefit of producing diverse solutions by taking advantage of the varied perspectives in our team. Getting the members of our team to sketch together created alignment among them, overcoming differences in background and skills^15^.

## Designing with users

We invited users to our workshops to give them the opportunity to contribute to the design activities. For example, we sketched together with users ways to show genetics data supporting a particular target-disease association (Fig. 2a), a data type that was identified as particularly important in one of our scenarios (Supplementary Fig. 4). Figure 2b shows the sketches that we created after two rounds of sketching. These sketches formed the basis for the visualization of mutations that we developed for the Platform (Supplementary Fig. 1d).

**Figure 2:**
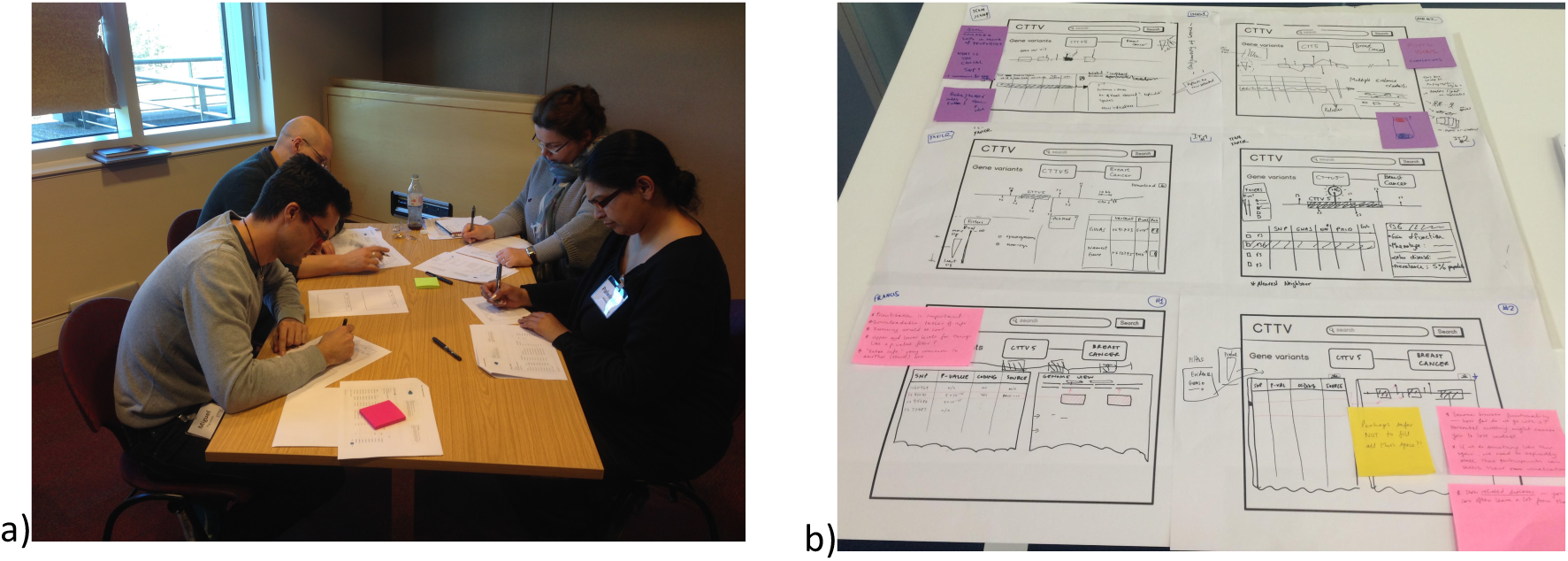
a) We sketched together with users ways to show genetics data supporting a particular target-disease association. b) The sketches that we produced after two rounds of sketching formed the basis for the visualization of mutations that we developed for the Platform (Supplementary Fig. 1d).

Engaging with users in this way helped us understand that they needed to be able to easily identify the mutations which are associated with a disease in the genomic context alongside other mutations. Clearly identifying this information is not always easy with more general genome browsing tools (such as Ensembl^16^ and UCSC^17^), which drug discovery scientists found overwhelming because of the volume of different data that they display.

The value of the design workshops was not restricted to their core outputs, i.e. the sketches that we created. Preparing the problem statements and identifying representative data samples for these workshops nurtured closer collaboration between the people working on the front end of the Platform and those who are responsible for data collection and processing.

The time we spent with users during the workshops was the most valuable outcome. This helped us further validate whether the questions that we were exploring with them were the right ones in the first place. Designing with users also made us more confident that the design will be in line with users’ expectations.

We ran three workshops with users focused on the overall Platform architecture and particular data sources in the period prior to the release of Beta v01. We continued to engage users in that way for improvements in subsequent versions of the Platform.

## Ensuring an intuitive solution through testing

The design phase is only the beginning of a search for a solution. Getting feedback about these designs from users (including people who were not involved in the sketching workshops) through user testing was critical to ensure that the Platform is intuitive.

In the period leading to the release of Beta v01 (Supplementary Fig. 7), we focused on collecting feedback from three to five users per iteration, engaging with each person for up to an hour^18^. Getting feedback very often from a small number of users has been shown to be more useful than larger user tests which may involve more participants but take place less frequently^19^.

For each testing session we prepared a guide which describes the problem that the design aims to solve and outlined the protocol that we would use to ask users for feedback. The developer responsible for implementing the design attended the session to observe users’ reactions first hand and ask questions. The feedback from these sessions was reviewed with whole team to identify the most important issues from each testing round.

Supplementary Figure 9 shows the iterative evolution of the Evidence page of the Platform from the initial sketch into versions of higher fidelity. This page presents the various kinds of evidence that associate a target with a disease, thus addressing one of Pat’s pain points (see Fig. 1). Each design in Supplementary Figure 9 is annotated with the feedback that we collected during the testing sessions.

Beta v01 was tried out by 20 target identification scientists from GlaxoSmithKline (GSK) who used it as part of their daily workflow. In addition to feedback sessions with each person, we organized focus group meetings with five to ten participants and at least one other member of the development team. The participants also sent us feedback by email. For example, we revised the default behaviour of the data type filters based on feedback that we received during this stage (see Supplementary Fig. 10).

After two iterations, we increased the number of Beta testers to 50, and after three more iterations to 100. Most of the feedback at these stages was about missing target-disease associations and incorrect assignments of supporting evidence. Given the wide range of domain expertise in our cohort of Beta testers this feedback was invaluable for creating a Platform that is trustworthy and comprehensive as well as intuitive.

Overall, we updated the Beta seven times (about once a month) based on the feedback that we collected from the Beta testers before making the Platform available to all drug target identification scientists at GSK R&D, the Wellcome Trust Sanger Institute and EMBL-EBI (the three founding partners of Open Targets). We revised the Platform again to address some outstanding issues that arose from this internal release and then made it publically available in December 2015.

Since the public release we had over 40 further engagements with more than 150 users in collaborative design activities and feedback sessions. We view developing the Platform as a process of continuous improvement and continue to strengthen it based on our regular interactions with users.

## Qualitative feedback and metrics

Qualitative feedback is at the heart of our iterative design process. In addition to helping us understand what needs to be improved, the feedback from users helps us assess whether we have met our main objective.

By the time that the Platform was made available publically we had collected plenty of feedback from users testifying that the Platform was comprehensive and intuitive. Users also shared examples of the Platform helping them with their day to day activities (see Supplementary Table 1).

We supplemented the qualitative feedback with quantitative metrics. These metrics were also defined collaboratively. Brainstorming a long list of metrics can quickly get unwieldy and hard to prioritize. To avoid this, we identified key performance indicators using the HEART methodology which breaks down the experience of using a product into five aspects: Happiness, Engagement, Adoption, Retention and Task completion^20^.

The importance of each of these aspects can vary depending on the product. We decided to focus on Adoption, Engagement and Retention (in that priority) for the definition of quantitative metrics, all of which can be captured regularly and more directly through web analytics. Task completion is less relevant to our application since using the Platform is much more open-ended than, for instance, making a purchase online (which has a clear completion action). Although we use qualitative feedback (Supplementary Table 1) as the main indicator of Happiness, we intend to start surveying our users periodically in order to monitor differences in the Net Promoter Score^21^ between major updates of the Platform.

We defined high level goals and lower level signals for the prioritized aspects as well as actual metrics for each aspect (See Supplementary Table 2). This process helped us think deeply and achieve clarity about the purpose of collecting analytics before investing effort in the actual way in this will be done.

When communicating our metrics to external stakeholders, we found that it was easier for them to comprehend if we replace the goals and signals column of Supplementary Table 2 with a simple question that the metrics are designed to answer as shown in Table 1. Table 1 reports the averages of these metrics from the beginning of April 2016 until the end of March 2017. The metrics suggest that the Platform has been used substantially during that year. This accords with the positive feedback that we have been receiving from users (Supplementary Table 1) and the fact that a new industrial partner (Biogen) joined Open Targets and started contributing to the development of the Platform soon after its public release.

**Table 1:** Key metrics for the period from the beginning of April 2016 until the end of March 2017. These metrics suggest that the Platform is being used substantially, in accordance to the positive feedback that we have been receiving from users.

|  | Question | Metrics | Average |
| --- | --- | --- | --- |
| Adoption | Are people visiting the site and viewing its pages? | Visits per week<br>Unique visitors per week<br>Pageviews per week<br>Unique pageviews per week | 816.35<br>529.58<br>4785.19<br>3343.42 |
| Engagement | Are people using the site and performing certain actions (internal site searches, downloads, clicking on evidence links)? | Average visit duration<br>Bounce rate per week<br>Actions per week<br>Actions per visit | 6min 49sec<br>25.60%<br>9401.33<br>11.29 |
| Retention | Are people coming back to the site? | Returning visits per week<br>% returning visits / all visits | 460.38<br>56.67% |

## Conclusion

There is little guidance on how to develop an intuitive web application for bench scientists. We have outlined how we designed such a web application for such scientists working on drug discovery. We hope that this will inform and inspire other members of the bioinformatics community wishing to design and develop more user friendly scientific web resources.

One common misconception about engaging with users is that it is restricted to asking them what they want, often after the application has been developed. We have demonstrated how we started engaging with users very early through user research and how we included them in collaborative design activities and regular feedback sessions within an iterative development process.

We applied Lean UX design methods to understand and meet the needs of our users. This approach is lean because we focused on understanding users’ needs before we started developing the application. However, the research, design and evaluation activities that we carried out produced very rich information about our users. This helped us empathize with them and move away from the ‘self-as-user’ outlook. By investing in user research, problem definition, design workshops and feedback sessions, we were more confident that we would deliver a useful and intuitive scientific application when it was made publically available.

We have also shown that developing our web application is a process of continuous improvement and have discussed how we supplement qualitative user feedback with quantitative metrics, which were also defined collaboratively.

## Author contributions

NK prepared and facilitated the reported UX activities and wrote the manuscript. FR and JAC led the user interviews. FR, JAC, D-CS, LF, SH, DH, GK, MM, WN, CKO, EP, AP, MP, SP, JV, XW JCB and ID contributed to the reported UX activities.

## Acknowledgements

We acknowledge funding from Open Targets, a pre-competitive partnership between Biogen, the European Bioinformatics Institute (EMBL-EBI), GlaxoSmithKline (GSK) and the Wellcome Trust Sanger Institute (WTSI). We are grateful to the participants of our studies, to Ewan Birney, Jonathan Hickford and Brendan Vaughan for their support and to Holly Foster for administrational assistance.

